# PsiPartition: Improved Site Partitioning for Genomic Data by Parameterized Sorting Indices and Bayesian Optimization

**DOI:** 10.1101/2024.04.03.588030

**Authors:** Shijie Xu, Akira Onoda

**Affiliations:** Graduate School of Environmental Science, Hokkaido University, Sapporo, 060-0810, Hokkaido, Japan; Faculty of Environmental Earth Science, Hokkaido University, Sapporo, 060-0810, Hokkaido, Japan

**Keywords:** Phylogenetics, Phylogenomics, Bayesian optimization, Partitioned models

## Abstract

Phylogenetics has been widely used in molecular biology to infer the evolutionary relationships among species. With the rapid development of sequencing technology, genomic data with thousands of sites becomes increasingly common in phylogenetic analysis, while heterogeneity among sites arises as one of the major challenges. A single homogeneous model is not sufficient to describe the evolution of all sites and partitioned models are often employed to model the evolution of heterogeneous sites by partitioning them into distinct groups and utilizing distinct evolutionary models for each group. It is crucial to determine the best partitioning, which greatly affects the reconstruction correctness of phylogeny. However, the best partitioning is usually intractable to obtain in practice. Traditional partitioning methods rely on heuristic algorithms or greedy search to determine the best ones in their solution space, are usually time-consuming, and with no guarantee of optimality. In this study, we propose a novel partitioning approach, termed PsiPartition, based on the parameterized sorting indices of sites and Bayesian optimization. We apply our method to empirical data sets and it performs significantly better compared to existing methods, in terms of Bayesian information criterion (BIC) and the corrected Akaike information criterion (AICc). We test PsiPartition on the simulated data sets with different site heterogeneity, alignment lengths, and number of loci. It is demonstrated that PsiPartition evidently and stably outperforms other methods in terms of the Robinson-Foulds (RF) distance between the true simulated trees and the reconstructed trees, especially on the data with more site heterogeneity. More importantly, our proposed Bayesian optimization-based method, for the first time, provides a new general framework to efficiently determine the optimal number of partitions. The corresponding reproducible source code and data are available at http://github.com/xu-shi-jie/PsiPartition.

## 1. Introduction

Phylogenetics provides powerful tools to reconstruct the evolutionary history of organisms based on their genetic sequences. In the past decades, various methods have been proposed to reconstruct phylogenetic trees such as the neighbor joining [1], maximum-likelihood (ML) [2], Bayesian inference (BI) [3], and graph splitting [4]. ML and BI are generally considered as the most accurate phylogenetic methods by modeling the evolution process as a continuous-time Markov process with a specific substitution model [5]. With the rapid development of sequencing technology, the size of the data sets has been increasing dramatically. Phylogenetics on genomic data, named phylogenomics, has become a hot topic in recent years. For example, DNA sequences from distinct genes of the species are often selected and concatenated into a supermatrix, to infer the phylogeny of species more accurately [6]. It has been suggested phylogeny inferred based on more sites is better than fewer [7]. However, the heterogeneity among sites becomes one of the major challenges in phylogenomic analysis. Genes in different regions of the genome may evolve at different rates, e.g., the third codon position of protein-coding genes evolves faster than the first and second codon positions [8] because the third codon position is often under less selective pressure. In addition, the heterogeneity of amino acids may arise from the different secondary structures of proteins [9]. Such heterogeneity is ubiquitous in genomic data and a single homogeneous model is not sufficient to describe it. This limits the potential improvements in the accuracy of phylogenetic inference by enhancing evolutionary signals with multiple genes [10]. Alternative methods that reduce the influence of site heterogeneity among sites are thus needed.

Partitioned models are proposed to describe the site heterogeneity and improve the accuracy of phylogenetic inference. In partitioned models, sites are divided into several disjoint subsets that apply different models, each of which may have different substitution rates, base frequencies, and branch lengths [11]. The importance of partitioned models has been demonstrated in phylogenomic analysis [6], as they can affect tree branches, topology, bootstrap support, and divergence date [12–17]. Many parameters of partitioned models affect the accuracy of phylogenetic inference and one of the most essential factors is the number of partitions. In general, more partitions can fit the data better as they utilize more evolutionary models. However, they also increase the risk of overfitting, which biases the subsequent analysis [18]. It is crucial to balance the goodness of fit and the complexity of the models. Information criterion such as the Akaike information criterion (AIC) [19], corrected AIC (AICc) [20], and Bayesian information criterion (BIC) [21] are often employed as an optimized objective in partitioned models.

There remain critical problems in partitioning sites efficiently and properly. Firstly, the number of all possible partitioning of an *n*-sites alignment is the Bell number *B*_*n*_satisfying the recursive definition 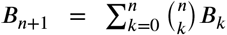 which increases faster than an exponential function as *n* increases. It is impossible to perform a brute-force search to exhaust all partitioning schemes even for alignments with a small number of sites [22]. Secondly, partitioning methods using *prior* knowledge such as codon positions and secondary structures of proteins are based on intuition, do not guarantee a correct partitioning of sites, and have therefore been improved in a series of works [12, 23–25]. Their methods partition sites based on a handful selection of collections of partitioning candidates, which requires expert knowledge and may lead to bias. Automatic partitioning methods have been proposed to search for a good partition by heuristic algorithms in the whole partitioning space. For example, the reported work [18] proposed a top-down greedy algorithm that partitions the sites from a single partition into more partitions gradually. It is based on the *k*-means clustering [26] of the site rates estimated from the reconstructed phylogenetic trees. Instead of partitioning sites, the reported work [27] recursively merges the most similar site pairs. These methods have achieved excellent performance while still having some disadvantages: 1) To acquire the estimated site rates, they rely on reconstructed reference phylogenetic trees, which take time to obtain and are possibly incorrect, and 2) they are based on greedy search and could fall in the local optimum. These disadvantages limit their applications in the situation with genomic data.

Recently, TIGER (Tree Independent Generation of Evolutionary Rates) has been proposed to estimate the evolutionary rates of sites without referring to phylogenetic trees [28]. It was originally developed to exclude the fast-evolving sites that interfere with the accurate reconstruction of actual phylogeny. TIGER-based partitioned models have demonstrated their success in many phylogenetic studies [29–37]. Rota *et al*. attempted to improve TIGER by using an extra “division factor” defined by users [38]. Kim *et al*. made efforts to automatically select the best number of partitions by recursive trisection [17]. These two methods alleviate the demands of automatic partitioning for genomic data. However, current TIGER-based methods still do not achieve the best accuracy. More accurate and efficient partitioning algorithms are thus needed.

In this study, we propose a novel site partitioning approach, termed PsiPartition, based on <u>P</u>arameterized <u>S</u>orting Indices (PSI) and Bayesian optimization. We first introduce the PSI as a better alternative to TIGER. Then we optimize PSI and the number of partitions by Bayesian optimization. We demonstrate that our method significantly outperforms the existing methods on empirical data sets in terms of Bayesian information criterion and corrected Akaike information criterion. Furthermore, it reconstructs the most accurate phylogenetic trees on the simulated data sets, in terms of Robinson-Foulds distance. More importantly, our method provides a new general Bayesian optimization-based framework to efficiently determine the best number of partitions and can be easily extended to other partitioning methods.

## 2. Method

### 2.1. Data sets

For phylogenetic inference benchmarking, we employ eight empirical DNA data sets from the reported work [16, 32, 33, 39–43]. Their statistics are listed in Table 1. These data sets are also employed as the benchmark data sets in previous studies [17, 38]. Alignments in the data sets have a range of base pair lengths from 4435 to 6372 and the number of taxa from 31 to 164. Each sequence in the alignments is the concatenated sequences from the mitochondrial gene (COI) and other four to seven nuclear genes commonly employed for lepidopteran phylogenetics, including CAD, EF-1 *α*, GAPDH, IDH, MDH, RpS5, and wingless [44].

**Table 1.** The empirical DNA data sets for benchmarking.

| Name | No. of taxa | No. of loci | No. of sites |
| --- | --- | --- | --- |
| Arctiina [39] | 113 | 8 | 5809 |
| Calisto [32] | 90 | 6 | 5297 |
| Choreutidae [16] | 41 | 8 | 6293 |
| Coenonymphina [40] | 69 | 5 | 4435 |
| Geometridae [41] | 164 | 8 | 5998 |
| Morpho [42] | 31 | 8 | 6372 |
| Noctuidae [43] | 78 | 8 | 6365 |
| Pieridae [33] | 110 | 8 | 6247 |

Six empirical protein data sets from the reported work [45] are also employed for benchmarking on the protein data. Their statistics are listed in Table 2. Alignments in the data sets have a range of amino acid lengths from 385 to 811 and the number of taxa from 17 to 31. Each alignment in the data sets includes the sequences of heat shock proteins Hsp83, Hsc70-3, Hsc70-4, Hsc70-5, Hsp60, and Hsp40. These proteins are the primary molecular chaperones protecting against thermal damage. Note that they are relatively small proteins and the number of sites is smaller compared to the DNA data sets.

**Table 2.** The empirical protein data sets for benchmarking.

| Name | No. of taxa | No. of sites |
| --- | --- | --- |
| Nguyen_2016a | 17 | 688 |
| Nguyen_2016b | 31 | 680 |
| Nguyen_2016c | 25 | 811 |
| Nguyen_2016d | 17 | 704 |
| Nguyen_2016e | 17 | 385 |
| Nguyen_2016f | 17 | 583 |

### 2.2. Parameterized Sorting Indices (PSI)

We introduce the proposed Parameterized Sorting Indices (PSI) by reviewing the Tree-Independent Generation of Evolutionary Rates (TIGER) [28]. In the original paper of TIGER, the evolutionary rates of site *i* are estimated as

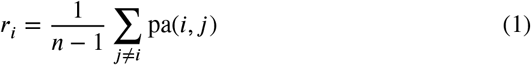

where *n* is the number of sites and

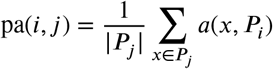

is the *agreement score*. Here the function *a*(*x, P*_*i*_) is defined as *a*(*x, P*_*i*_) = 1 if *x* ⊂ *A* and *A* ∈ *P*_*i*_, and otherwise 0. The set *P*_*i*_ is the *set partition* of the site *i*, whose elements are disjoint subsets of {1, 2, …, *m*} where *m* is the number of taxa. For each subset *A* ∈ *P*_*i*_, the character of the *l*-th sequence at site *i* has the same type for all *l* ∈ *A*. For example, consider two sites *S*_*i*_ = CTTAA and *S*_*j*_ = AGGGG with the set partition *P*_*i*_ = {{1}, {2, 3}, {4, 5}} and *P*_*j*_ = {{1}, {2, 3, 4, 5}}, respectively. We have pa(*A, B*) = 0.5 because out of two subsets {1} and {2, 3, 4, 5} ∈ *P*_*j*_, only {1} ⊂ {1} ∈ *P*_*i*_. pa(*B, A*) = 1 because for all subsets {1}, {2, 3}, {4, 5} ∈ *P*_*i*_, we have {1} ⊂ {1} ∈ *P*_*j*_, {2, 3} ⊂ {2, 3, 4, 5} ∈ *P*_*j*_, and {4, 5} ⊂ {2, 3, 4, 5} ∈ *P*_*j*_. The TIGER of site *S*_*i*_ and *S*_*j*_ should be *r*_*i*_ = 3*/*3 = 1 and *r*_*j*_ = 1*/*3, respectively.

PSI has two key innovations. Firstly, in the original TIGER, the function *a*(*x, P*_*i*_) is defined as binary values 1 or 0 for all characters *x* at site *j*, regardless of the characters’ types. However, characters representing different base pairs or amino acids should have different properties and should be treated differently. This is similar to the idea of base frequency parameters implemented in most phylogenetic inference tools [46, 47]. We therefore introduce a group of parameters *w* to represent these differences among characters. For example, we adopt five parameters *w* = {*w*_*A*_, *w*_*C*_, *w*_*G*_, *w*_*T*_, *w*_gap_} for DNA data sets, where *w*_*A*_ is the parameter for character *A, w*_gap_ is the parameter for character gaps or missing characters, and so on. We define *a*_*w*_(*x, P*_*i*_) = *w*_*p*(*x*)_ if *x* ⊂ *A* and *A* ∈ *P*_*i*_ and otherwise 0 where *p*(*x*) is the corresponding character of subset *x*. Similarly, there should be 21 parameters for protein data sets.

Secondly, instead of using Equation 1, we utilize a comparison between two sites to define PSI at site *i* by following

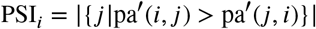

where pa′(*i, j*) is the defined as following

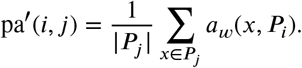

PSI_*i*_ ranging from 0 to *n −* 1 represents the number of sites that site *i* agrees more than it is agreed by.

After calculating the PSI for all sites, we bin the sites into *k* partitions based on their PSIs. Denote *α* = min_*i*_ PSI_*i*_ and *β* = max_*i*_ PSI_*i*_, then *k* partitions are defined as following

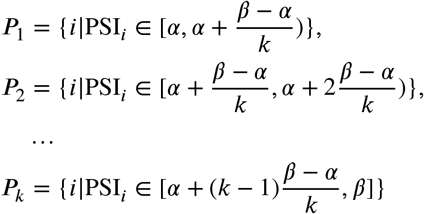

where *k* is the number of partitions. For example, if *α* = 0 and *β* = 10, then 3 partitions are 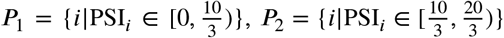, and 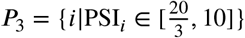.

Note that the parameters *w* and the number of partitions *k* described in this section are not determined and will be optimized based on the input data.

### 2.3. Bayesian optimization for PSI

It is important to decide the best number *k* of partitions in the partitioned models. When *k* equals the number of sites, a single-site partitioning can fit the data very well. This may cause overfitting and produce the reconstructed phylogeny different from the true one. When *k* = 1, the single partition is not suitable for describing heterogeneity among the sites. Some existing methods employ greedy algorithms to determine the number of partitions which is time-consuming for large data sets.

Bayesian optimization (BO), a black-box method, has been widely used in many fields for parameter tuning and are also adopted in this study. It has a siginificant advantage when the objective function is expensive to evaluate and the number of parameters is small. In our situation, the objective function is the information criterion (i.e., BIC) of the partitioned models, which is expensive to evaluate because it requires full phylogenetic inference on the given data. The optimized parameters are the number of partitions *k* and the introduced parameters *w* of PSI, which are few (i.e., 6 for DNA and 22 for proteins). BO utilizes a surrogate model such as the Gaussian process model [48] to approximate the objective function. The parameters are estimated based on the acquisition functions such as expectation improvement and upper confidence bound. Then BO evaluates the objective function on these parameters to obtain a new observation and updates the surrogate model with all observations. This process repeats until the parameter converges or the maximum number of iterations is reached.

We propose PsiPartition algorithms based on BO, as shown in Algorithm 1. The input of the algorithm is the alignment *A*, the maximum number of partitions *k*_max_, and the maximum iterations steps *n*. The output is the optimal partition *P*\*, the optimal parameters *s*\* and the optimal objective value *o*\*. The algorithm firstly initializes an empty set *S* and samples the number of partitions *k*_1_ from {2, 3, …, *k*_max_} and the parameters *w*_1_ from [0, 1]_*t*_ where *t* = 5 for DNA data and *t* = 21 for amino acid data. For each iteration *i*, it calculates the PSI *ψ*_*i*_ of all sites with parameters *w*_i_ and bins the sites into *k*_i_ partitions 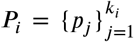 based on *ψ*_*i*_. It thus evaluates the phylogenetic software (e.g., IQ-TREE) with the partitioning *P*_*i*_ to obtain the BIC *b*_*i*_. Each evaluation is recorded as an observation (*P*_*i*_, *s*_*i*_, *b*_*i*_) where *s*_*i*_ = (*w*_*i*_, *k*_*i*_), and pushed into the set *S*. Then it updates the surrogate model *m* with all observations (*P*_*i*_, *s*_*i*_, *b*_*i*_) ∈ *S* and determines the next parameters *s*_*i*+1_ = (*w*_*i*+1_, *k*_*i*+1_) by the surrogate model *m* with expected improvement. The iterations will be repeated for *k*_max_ times. Finally, the algorithm returns the partitioning *P*\*, the parameters *s*\*, and the BIC *o*\* that has the minimum BIC *o*\* in *S*.

To indicate the necessity of BO, we develop a variant of PsiPartition, termed PsiPartitionFast, which freezes all parameters *w*_*i*_ as 1 in Algorithm 1. Compared to PsiPartition, it only optimizes the number of partitions *k*. The search space can be reduced and the overall steps of optimization are also less than PsiPartition. A detailed description is referred to the pseudocode in Supplementary Algorithm S1.

#### Algorithm 1 PsiPartition

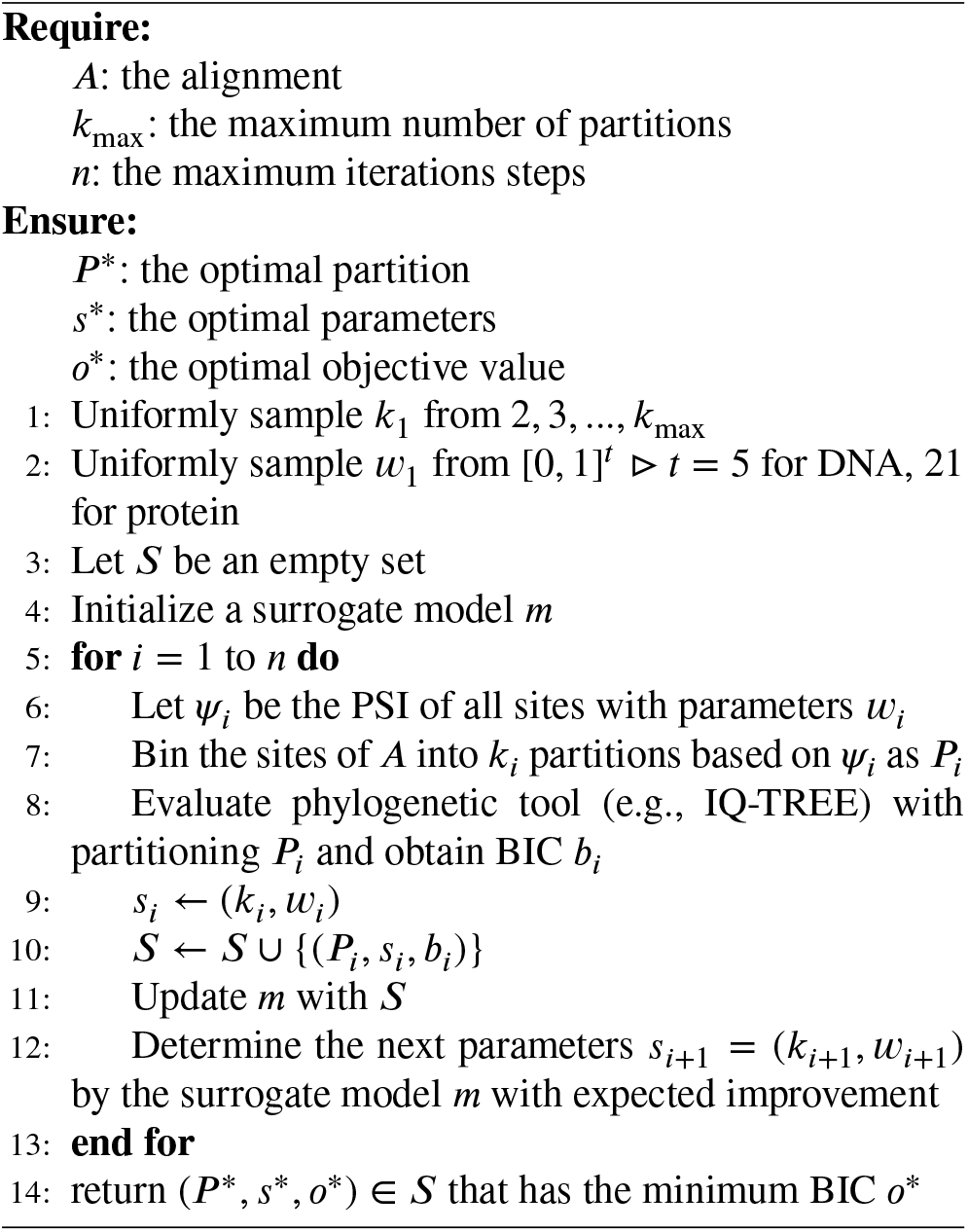

### 2.4. Evaluation metrics

To evaluate the performance of different partitioning methods, we employ the Bayesian information criterion (BIC) [21] and the corrected Akaike information criterion (AICc) [20]. The BIC and AICc are defined as the following

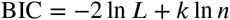

and

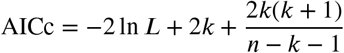

where *L* is the likelihood of the model, *k* is the number of model parameters, and *n* is the number of sites. BIC and AICc are both used to assess the goodness of partitioned models, which with the smallest BIC or AICc are considered the best ones. The BIC is the default criterion for model selection in IQ-TREE software [49]. We thus adopted it as the major evaluation metric and AICc as a reference. It is worth noting that the partitioning of sites is usually considered as *a priori*. Therefore the parameters *w* introduced in the aforementioned Bayesian optimization algorithm and the number of partition *k* should not be included in the calculation of BIC and AICc.

The Robinson-Foulds (RF) distance [50] is adopted to evaluate the accuracy of the reconstructed phylogenetic trees. Two branches sharing a common ancestor in a tree split a part of taxa into two disjoint subsets (i.e., clades). Therefore they recursively imply a nested partitioning of the taxa. The RF distance is defined as the summation of the number of partitions implied by one tree but not by another one. It is a metric of the topological distance between two trees, and the smaller the RF distance, the more similar the two trees. In our experiments, we compute the RF distance between the true simulated trees and the reconstructed trees and compare them of different partitioning methods.

We utilize the transfer distance to compare the similarity between two partitions. Suppose *X* be a set, and *P* = {*C*_1_, *C*_2_, …, *C*_*n*_} and 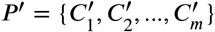 are two partitions of *X* Without loss of generality, we assume *n* = *m* because we can add empty subsets to the smaller one. We firstly define the *concordance* as following

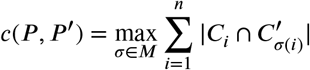

where *M* is the set of all permutations of {1, 2, …, *n*}. The *transfer distance* is defined as following

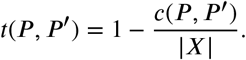

It is obvious to see that 0 *≤ t*(*P, P* ′) *≤* 1 and *t*(*P, P* ′) = 0 if and only if *P* = *P* ′. A smaller transfer distance means more similarity between two partitions. The proof of the properties of the transfer distance is described in [51] with more details.

## 3. Results

PsiPartition is implemented within Python 3.12.1. To perform the Bayesian optimization, wandb package (version 0.16.1) [52] are also used [53]. It employed normalized Gaussian process model with Matern kernel [54] and parameter *α* = 10^*−*7^. Expected improvement is utilized as the default acquisition function. The parameters *k*_max_ and *n* in Algorithm 1 are both set up to 30 by default unless mentioned. The partitioned models and their information criterion are evaluated by ModelFinder [49] implemented in IQ-TREE [46] (version 1.6.12). UFBoot [55] in IQ-TREE is also enabled for ultrafast bootstrap calculation of phylogenetic trees. The experiments are performed on a Linux machine with an AMD Ryzen 9 7950X3D 16-core processor and 64 GB memory. Multiple threads feature of IQ-TREE are adopted to accelerate the phylogenetic inference. Interactive Tree Of Life (ITOL) online tools [56] are used to visualize the phylogenetic trees and the bootstrap support values of branches.

### 3.1. PsiPartition outperforms other partitioning methods on empirical data sets

To evaluate the performance of our proposed methods, PsiPartition and PsiPartitionFast, and existing state-of-the-art methods, we applied them to eight empirical DNA data sets listed in Table 1. The experimental results are shown in Fig. 1. The compared methods include the default IQ-TREE using the single homogeneous model without partitioning (NP), RatePartition with parameter `div_factor` = 4 (RatePartition-4) and `div_factor` = 5 (RatePartition-5), and mPartition. We show the advantages of partitioned models compared to NP by BIC decreases defined as the following ΔBIC = BIC of partitioned models *−* BIC of model without partitioning, and ΔAICc is defined similarly.

**Fig. 1.**
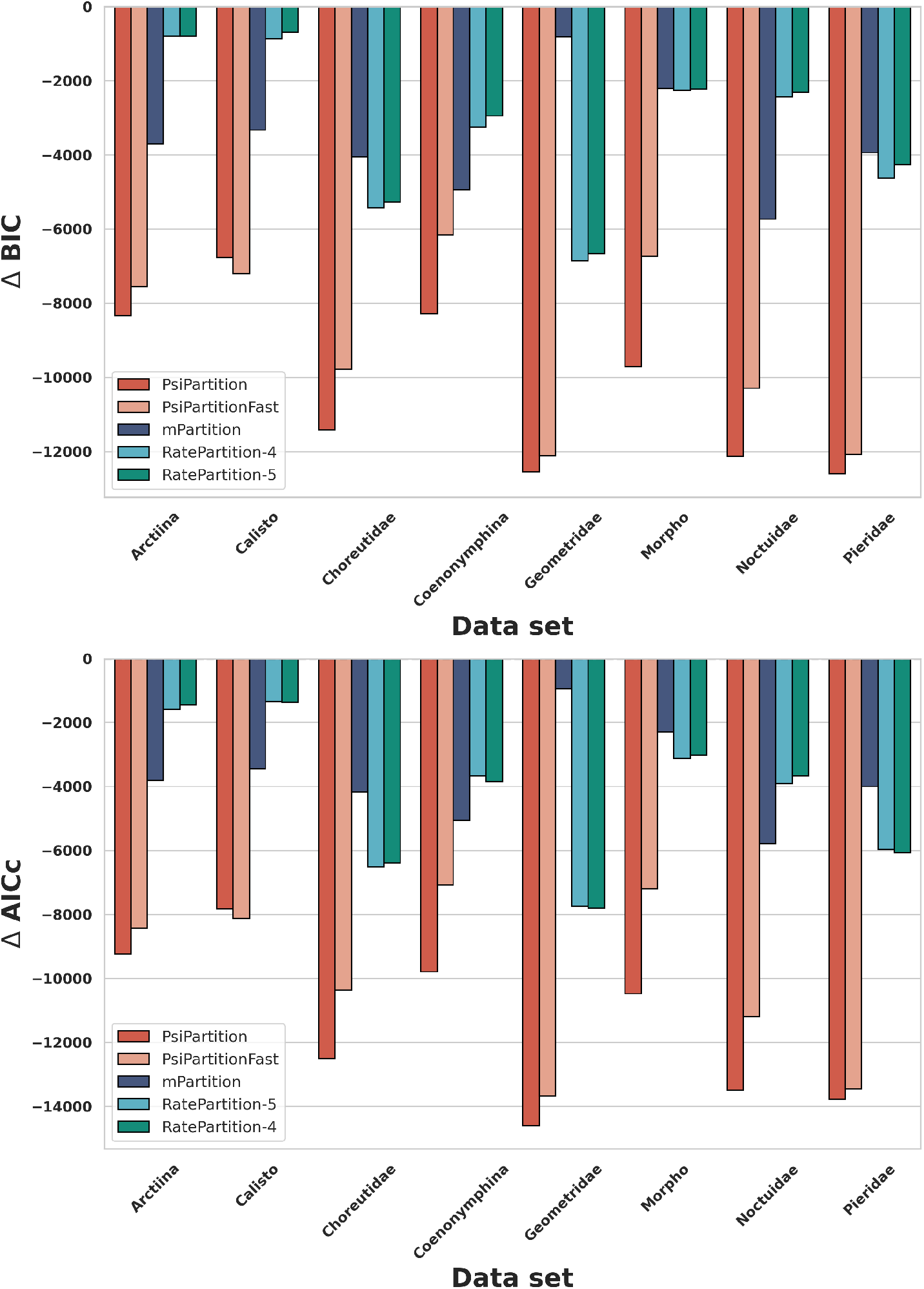
The performance comparison of different partitioning methods on the empirical DNA data sets (Arctiina, Calisto, Choreutidae, Coenonymphina, Geometridae, Morpho, Noctuidae, and Pieridae). Our proposed methods (PsiPartition and PsiPartitionFast) achieves the best performance in terms of the ΔBIC and ΔAIC, significantly outperforming the other methods.

On all data sets, our proposed methods PsiPartition and PsiPartitionFast significantly outperform other partitioned methods in terms of the ΔBIC and ΔAICc. Both of two significantly reduce the BIC and AICc values of the partitioned models, as shown in Supplementary Table S1. In addition, PsiPartition is better than PsiPartitionFast, which suggests that the parameters *w* play a vital role for partitioning. On the data set Calisto, PsiPartitionFast is slightly better than PsiPartition, which is probably caused by the relatively small size of the data set and the randomness in phylogenetic inference [57]. The results demonstrate that our proposed methods are more accurate than the default IQ-TREE and other existing partitioning methods, especially on the data sets with more sites and taxa.

We visualize the distribution of partitions in Fig. 2. Considering the differences among data, the number of partitions *k* determined by PsiPartition for different empirical DNA data sets are distinct, ranging from 13 to 30. This suggests that our proposed methods are capable of searching for different optimal partition schemes on different data sets. Furthermore, the number of sites that fall into each partition are also plotted. Bin indices represent the binning of sites based on the PSI, i.e., the sites with lower PSI will be assigned to the smaller bin indices, and vice versa. It is clear that the distribution of the number of sites in each bin is quite imbalanced and most of the sites are assigned to the first several bins, while there are still a few sites assigned to the last several bins. This is similar to those in previous study, suggesting that our methods properly partition the sites.

**Fig. 2.**
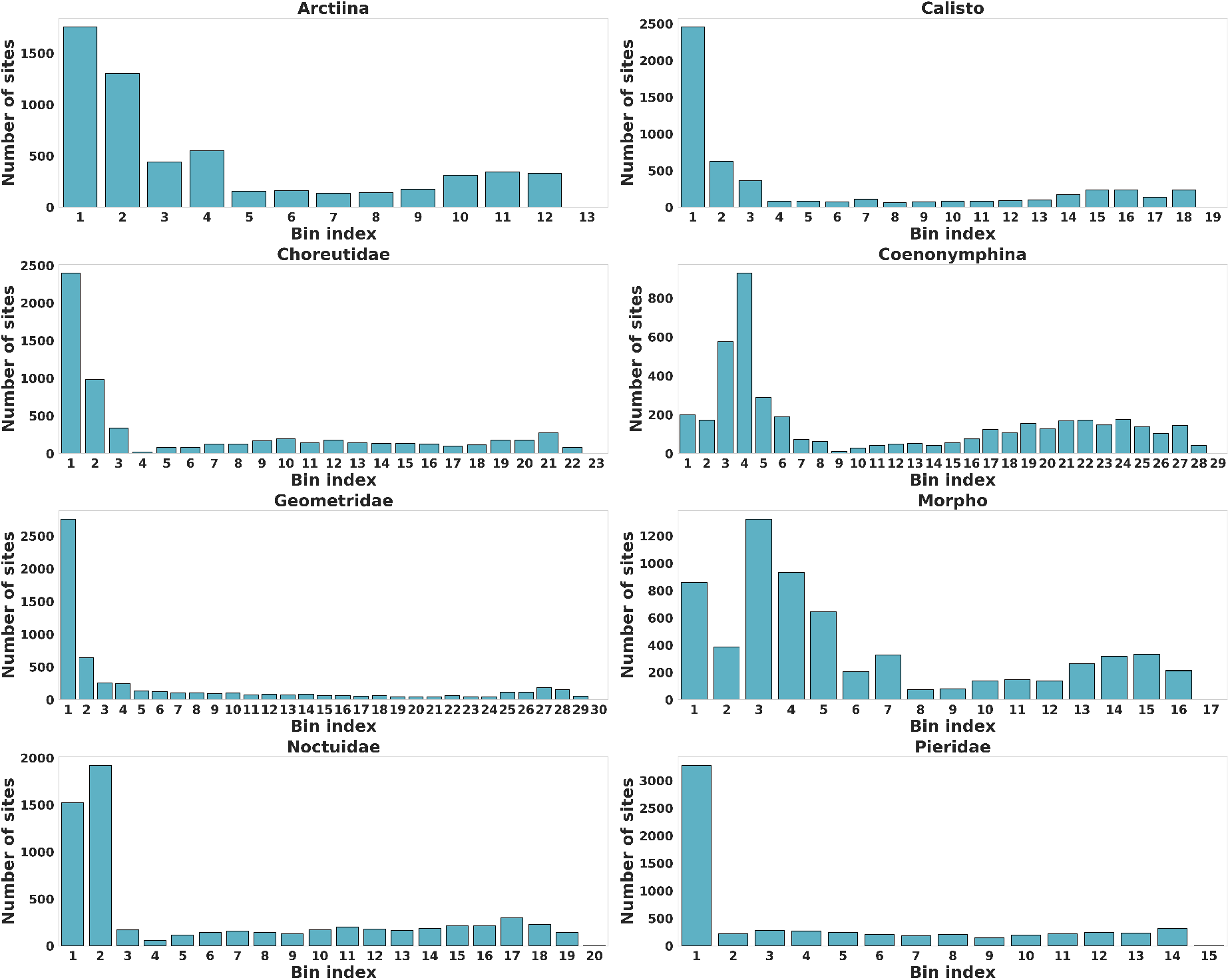
The number of sites in each bin generated by PsiPartition on the different empirical DNA data sets (Arctiina, Calisto, Choreutidae, Coenonymphina, Geometridae, Morpho, Noctuidae, and Pieridae). Different bins have different number of sites, with a long tails of bins with small number of sites. Note here a smaller bin index means a smaller PSI compared to the other bins, suggesting lower estimated evolutionary rates.

PsiPartition and PsiPartitionFast produce totally different partitioning compared to existing methods. The transfer distances between the partitions generated by our proposed methods and other existing methods are shown in Fig. 3. Here the X-axis and Y-axis represent the partitions generated by different methods and blocks with smaller values and darker colors mean more similarities between the two partitions. The results demonstrate that the partitions generated by our proposed methods, PsiPartition and PsiPartitionFast, have quite large transfer distance to the partitions generated by remaining methods. On some data sets such as Morpho and Coenonymphina, the partitions generated by PsiPartition and PsiPartitionFast are also considerable different from each other. Interestingly, two existing methods, RatePartition-4 and RatePartition-5 generated quite similar partitions to each other, because they utilized identical TIGER with only different divsion factors. Our PsiPartition instead optimized the PSI separately even on the same data set. These results suggest that our proposed methods are capable of searching for different partition schemes effectively and potentially finding the optimal ones.

**Fig. 3.**
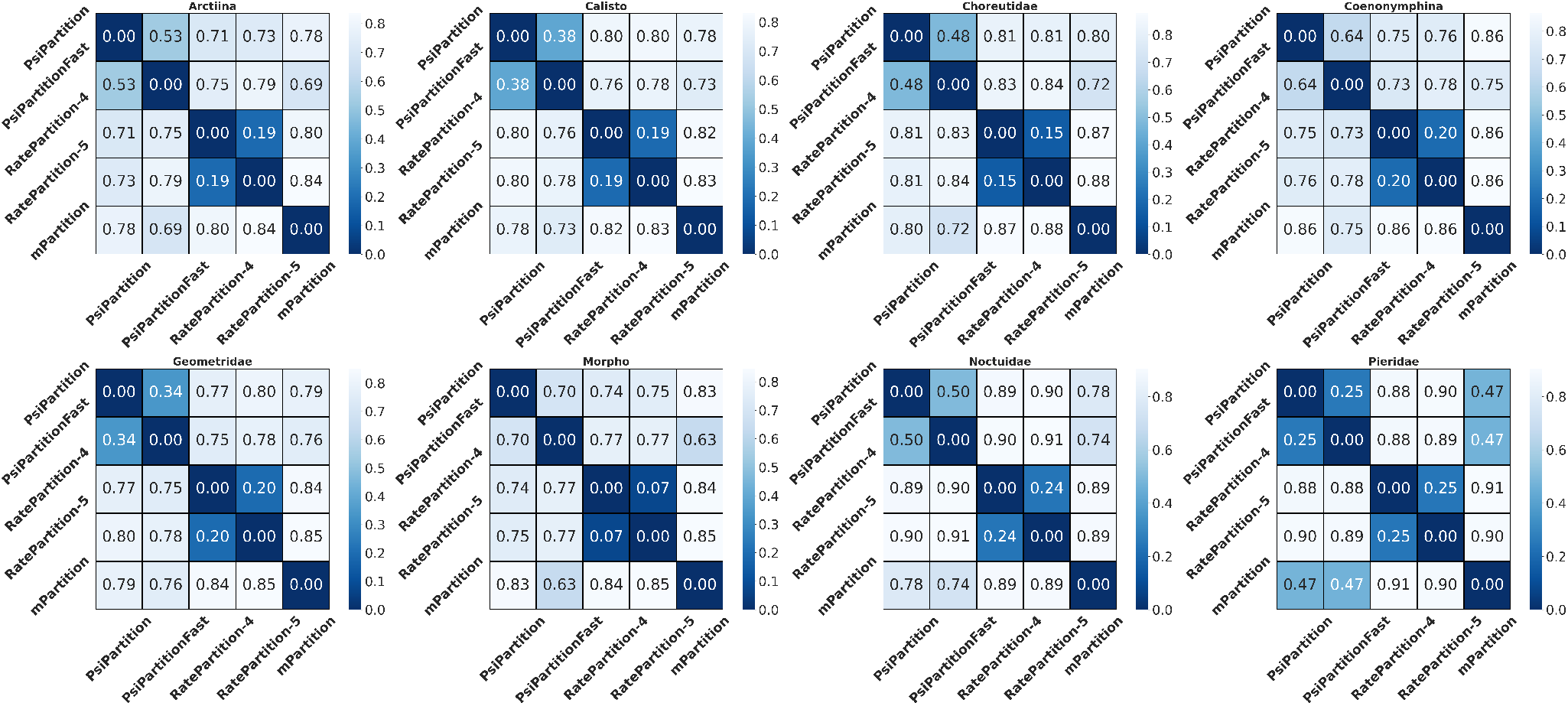
The transfer distances between the partitions generated by our proposed methods (PsiPartition and PsiPartitionFast) and other methods on different empirical DNA data sets (Arctiina, Calisto, Choreutidae, Coenonymphina, Geometridae, Morpho, Noctuidae, and Pieridae). PsiPartition and PsiPartitionFast have large transfer distances to the other methods, suggesting their partitions are significantly different from the other methods.

Our proposed methods are also superior on the protein data. We evaluate the performance comparison of different partitioning methods on the empirical protein data sets, as shown in Fig. 4. On most data sets, our proposed PsiPartition and PsiPartitionFast significantly reduce the BIC and AICc, as shown in Supplementary Table S2. PsiPartition significantly outperforms other methods in terms of the ΔBIC and ΔAICc. The only exception is on the data set “Ngyuen_2016e” where the PsiPartition is slightly worse than the RatePartition-4 and RatePartition-5. This is because the Ngyuen_2016e data set has a relatively small number of sites (385 amino acids) while others have about 600 sites or more. As the number of sites increases, the heterogeneity among sites could increase. The improvements achieved by our proposed methods compared to existing methods become more significant. Additionally, the PsiPartitionFast also achieved comparable performance to other methods. It is slightly worse than the PsiPartition because it does not adopt Bayesian optimization. The results demonstrate that our proposed PsiPartition is more accurate than the default IQ-TREE and other methods. By comparing PsiPartition and PsiPartitionFast, it is suggested that Bayesian optimization is necessary for accurate phylogenetic inference.

**Fig. 4.**
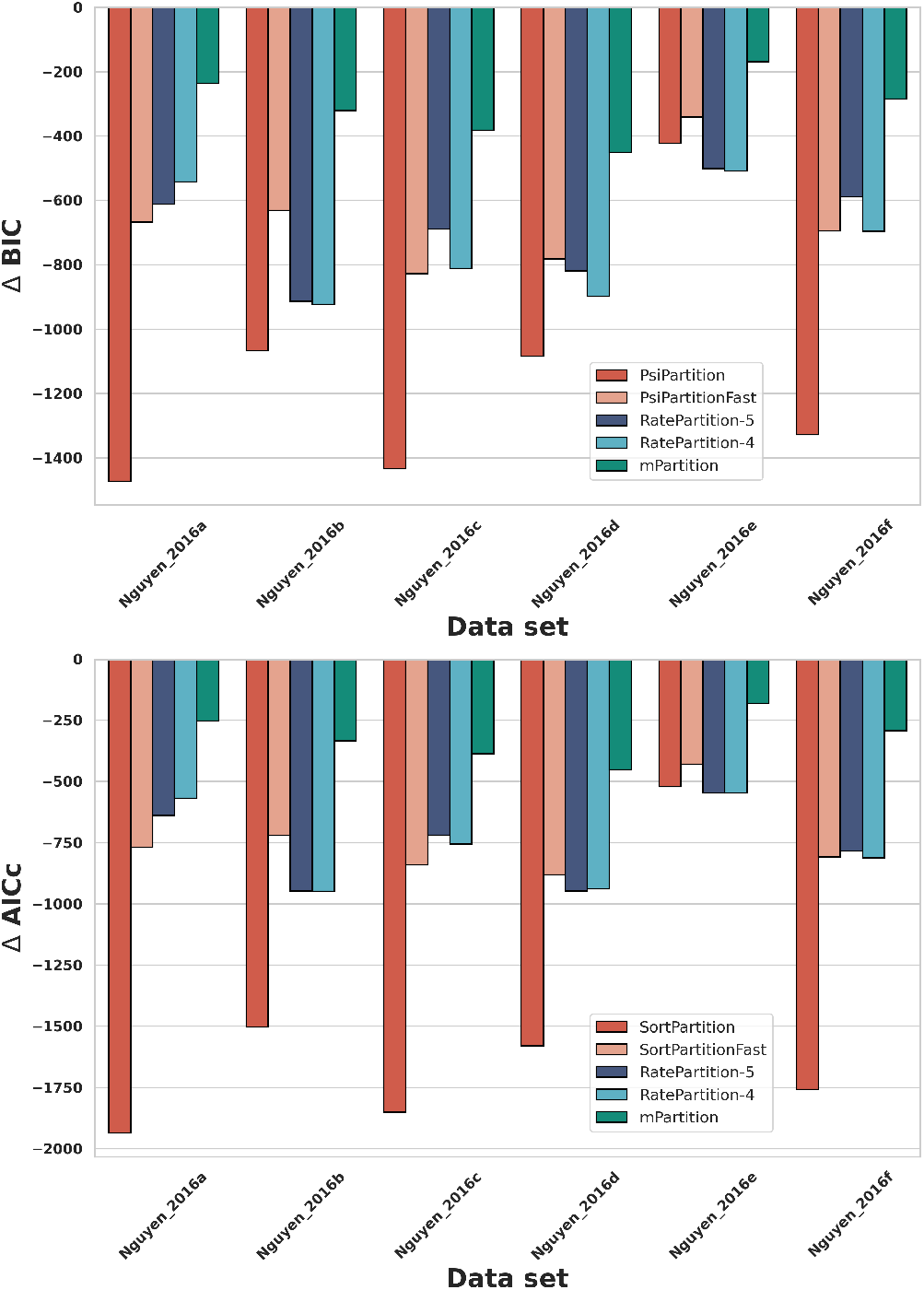
The performance comparison of different partitioning methods on the empirical protein data sets (Ngyuen_2016[a-f]). Our proposed method PsiPartition achieves the best performance in terms of the ΔBIC and ΔAIC values, significantly outperforming the other methods.

### 3.2. Bayesian optimization efficiently determines the optimal parameters of PsiPartition

We further assess the efficiency of PsiPartition on different empirical DNA data sets. The tracking of the evaluated BIC and AICc during the optimization iterations is shown in Fig. 5. The results demonstrate that the Bayesian optimization gradually decreases BIC and finally converges (or fluctuates) at the optimal BIC values. AICc can be similarly anlayzed. More specifically, we list the steps of iterations to achieve the best BIC and AICc in Table 3. These optimizations take a range of 8 to 29 steps (17.8 on average) to achieve an optimal BIC. Similarly, they take 9 to 29 steps (19.1 on average) to achieve an optimal AICc. The results illustrate that our proposed methods can expeditiously determine the optimal parameters in a limited steps of iterations.

**Table 3.** The numbers of iterations to achieve the best BIC and AICc on different empirical DNA data sets.

| Name | # Iter. to opt. BIC | # Iter. to opt. AICc |
| --- | --- | --- |
| Arctiina | 12 | 12 |
| Calisto | 8 | 9 |
| Choreutidae | 29 | 29 |
| Coenonymphina | 22 | 22 |
| Geometridae | 28 | 25 |
| Morpho | 16 | 23 |
| Noctuidae | 15 | 15 |
| Pieridae | 12 | 18 |

**Fig. 5.**
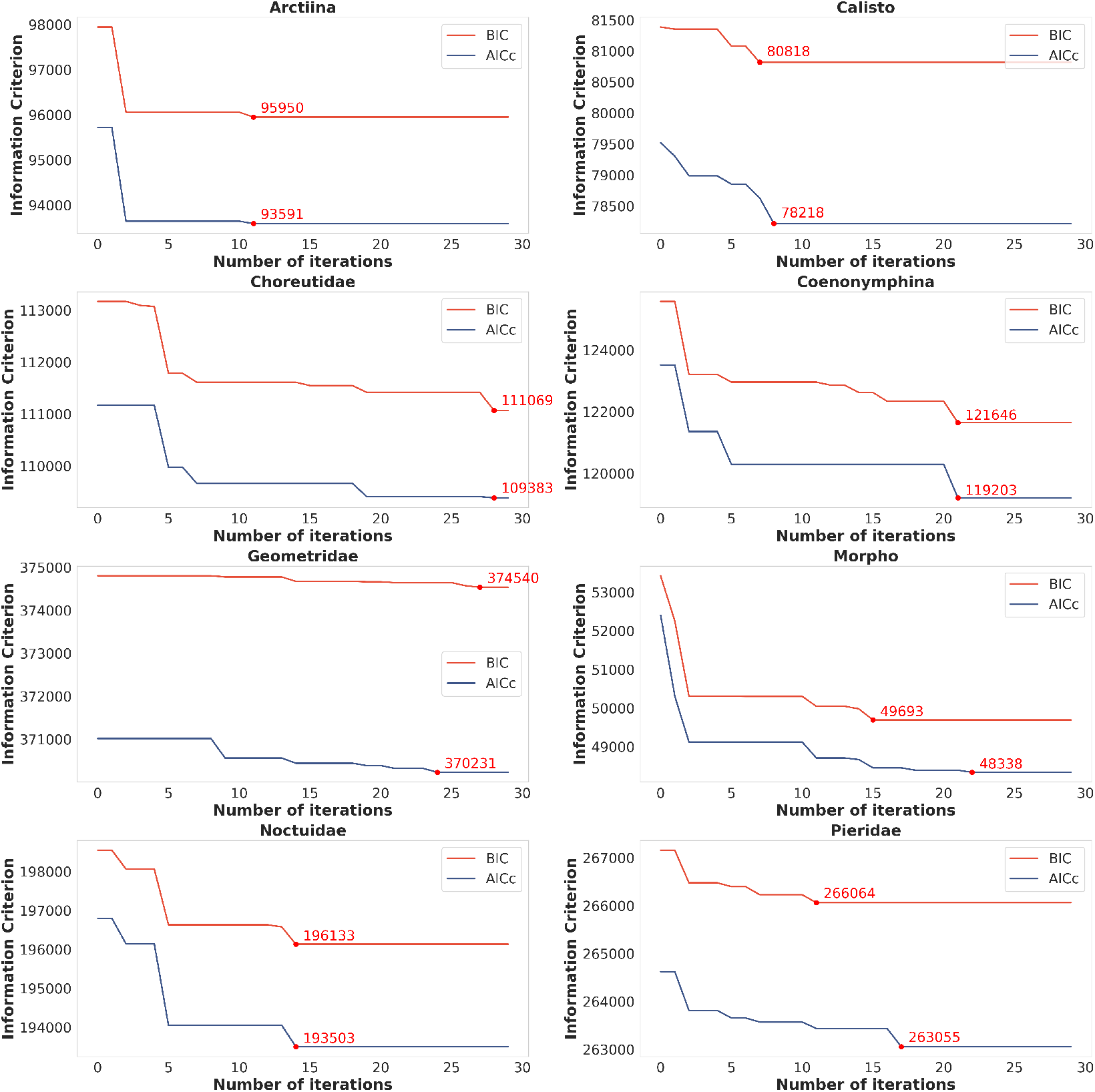
Bayesian optimization for different empirical DNA data sets.In the curves, each point (*k, v*) where *k* = 1, 2, …, 30 represent that PsiPartition achieved the minimum values of *v* of BIC (red) or AICc (red) in the first *k* iterations. The highlighted points with annotated values in the curves are the best information criterion values achieved by PsiPartition for the first time through the whole curves.

It is intractable to find the global minimum of BIC on all possible parameters, therefore we run PsiPartition 1000 times for the Morpho data set with uniformly sampled random parameters *w* from [0, 1]^*t*^ and *k* from {2, 3, …, *k*_max_}, respectively. We check whether Bayesian optimization-based minimum BIC achieves the best among all randomly evaluated BICs. Considering the number of parameters of PsiPartition on DNA data sets is six, t-distributed Stochastic Neighbor Embedding (t-SNE) [58] is employed to embed parameters into the 2-dimensional plane, which will keep the Euclidean distance between the parameters as much as possible. We normalize each parameter to normal distribution before applying t-SNE to make them comparable. The BIC values of each evaluation are shown as the sizes and colors of the spots (darker and larger spots mean smaller BIC, and vice versa). As shown in Fig. 6, it is clear that most of the evaluations from Bayesian optimization produce sufficiently good BIC values, which are concentrated into a cluster in the left part. The optimal parameters from Bayesian optimization (circled in orange) also achieve the best BIC among all random evaluations. These results suggest that PsiPartition is capable of approaching the best parameters efficiently compared to the random search.

**Fig. 6.**
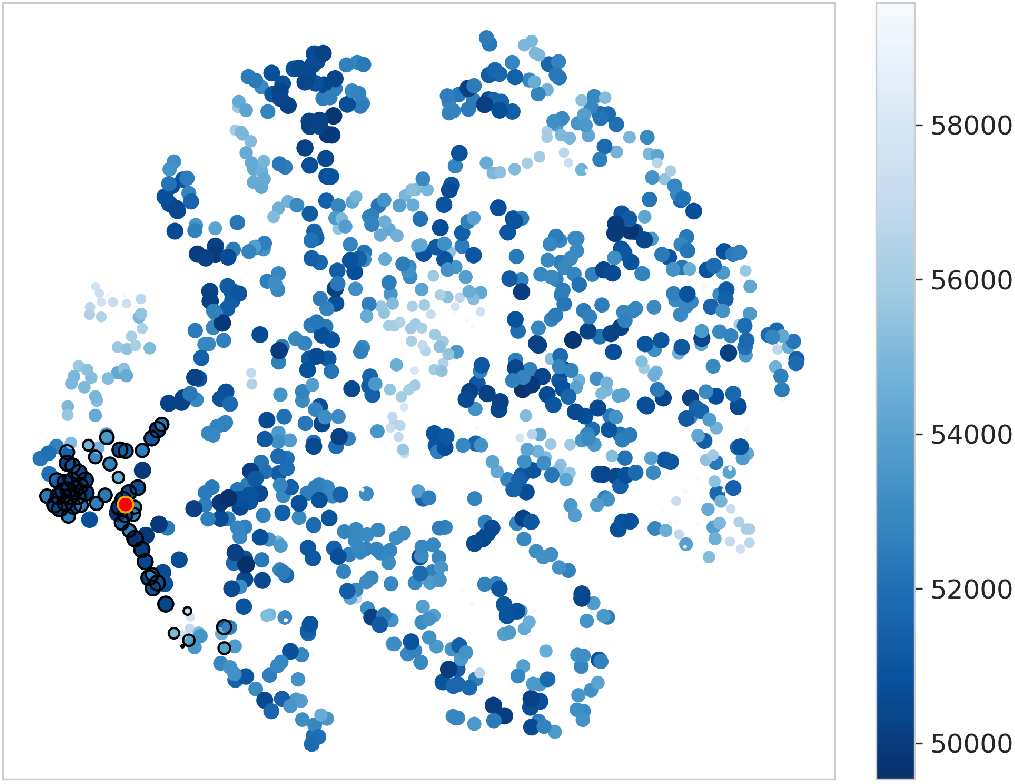
Scatter diagram of BICs produced by PsiPartition with different parameters *w* and *k* on the Morpho data set. The trajectory of Bayesian optimization is circled with black and the best Bayesian optimized BIC is the red spot circled with orange.

### 3.3. PsiPartition accurately reconstructs simulated phylogenetic trees

To evaluate the performance of our algorithm on simulated data sets, we generate 100 simulated DNA alignments with 49 sequences and 5000 sites. We employ Seq-Gen software [59] and the General Time Reversible (GTR) model [5] with 4-category Gamma distribution to generate the alignments from 100 randomly generated trees, which are described in Supplementary Algorithm S2. We utilize Seq-Gen’s parameters “`-r rate_matrix -f base_freq`”, where `rate_matrix` and `base_freq` are 6- and 4-dimensional vectors. Their elements are randomly generated from a uniform distribution *U*[0, 1] and represent the rate matrix and base frequencies, respectively. For `base_freq`, an additional *L*_1_ normalization is performed to make them indeed base frequencies. We segmented 5,000 sites into 1 to 10 equal-length loci and applied Seq-Gen to each locus then concatenated the produced alignments of each locus into the single 5000-site alignments. This process is repeated 10 times for each number of loci and finally, we obtained 100 simulated alignments. We denoted this data set as “SimDNA-1”. PsiPartition and PsiPartitionFast present a superior performance compared to other methods on the simulated data set SimDNA-1 in terms of the RF distance between the true trees and the reconstructed trees, as shown in Fig. 7. The standard deviations of the RF distances are also shown. The results demonstrate that, for genomic data with heterogeneous site rates, our methods are robust and more accurate than the single homogeneous model without partitioning and existing partitioning methods. We emphasize that the differences of performance between our methods and others on the data sets with only one locus are not significant while the differences become larger in the data sets with more loci. Such differences do not increase when the number of loci continues to increase because the total lengths of sequences are always 5,000. This suggests the necessity of partitioning the sites by PsiPartition for large data sets. It is worth noting that both RatePartition-4 and RatePartition-5 failed to run on some of the simulated data sets with 87% (87/100) success rate because some sites are incorrectly partitioned and only one state is observed in some partitions thus IQ-TREE failed to start phylogenetic inference.

**Fig. 7.**
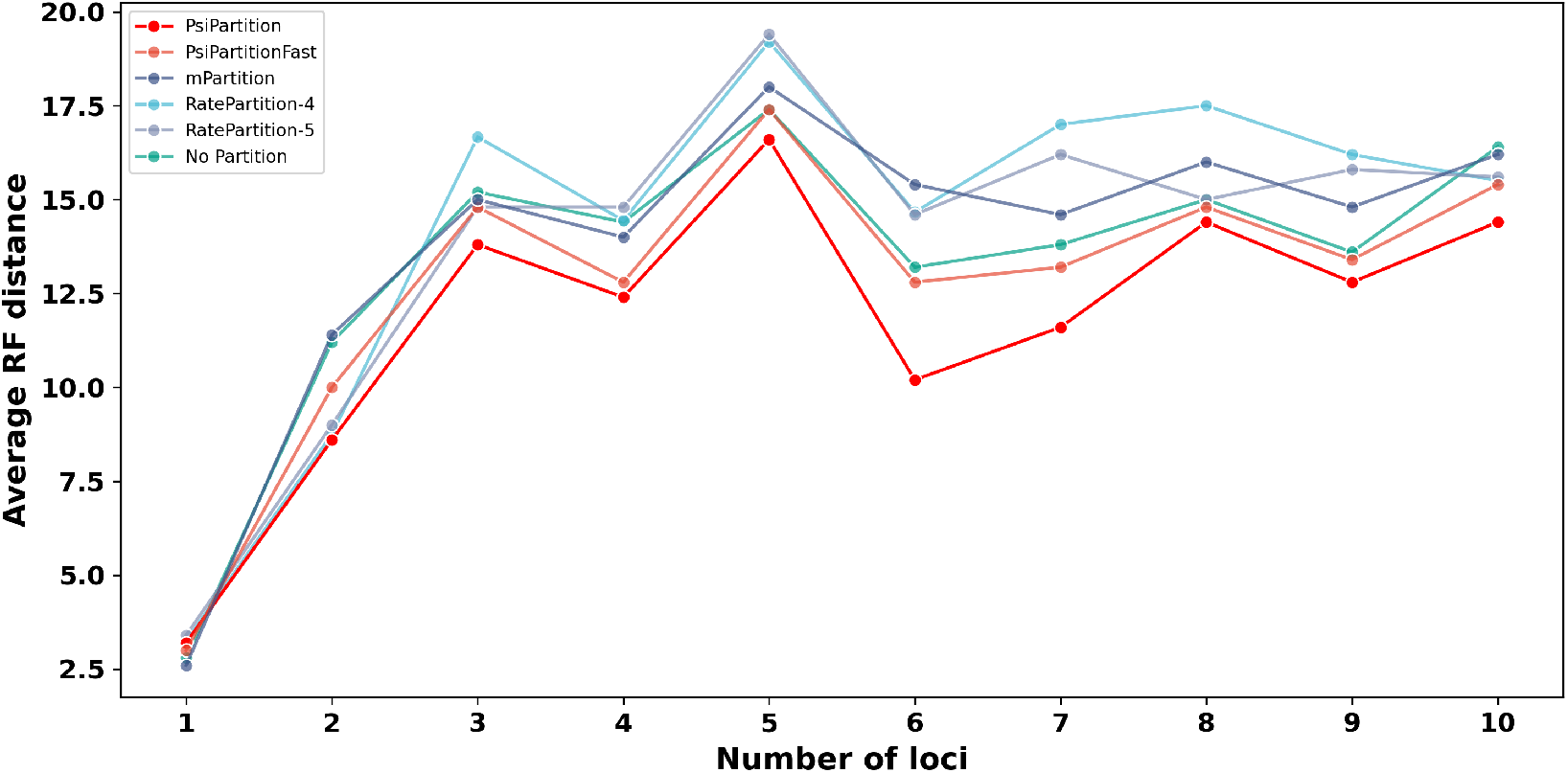
The average RF distances between the true trees and the reconstructed trees on SimDNA-1. For each different number of loci, our proposed PsiPartition (red) reconstructs the most accurate trees, with the smallest RF distances. The advantages of PsiPartition compared to other methods are significant when the number of loci is larger than 2.

We also generate 100 simulated DNA alignments with 6 to 15 loci and the lengths of single locus are always set up to 1,000. That is, we obtain alignments that have a range of lengths from 6,000 to 15,000 and each length has 10 alignments. The algorithms and configurations to generate them are the same as aforementioned. We denote this data set as “SimDNA-2”. We compare the performance of different methods in terms of RF distance, as shown in Fig. 8. It is clear that our PsiPartition significantly outperforms other methods on most of the alignment. The only exception when the number of loci is 10 can be explained as a random bias during the phylogenetics inference because only 10 alignments are evaluated for each length. It is worth noting that some methods cannot reconstruct all phylogenies on SimDNA-2 (mPartition reconstructs 99 and RatePartition-5 reconstructs 93 out of 100 phylogenies).

**Fig. 8.**
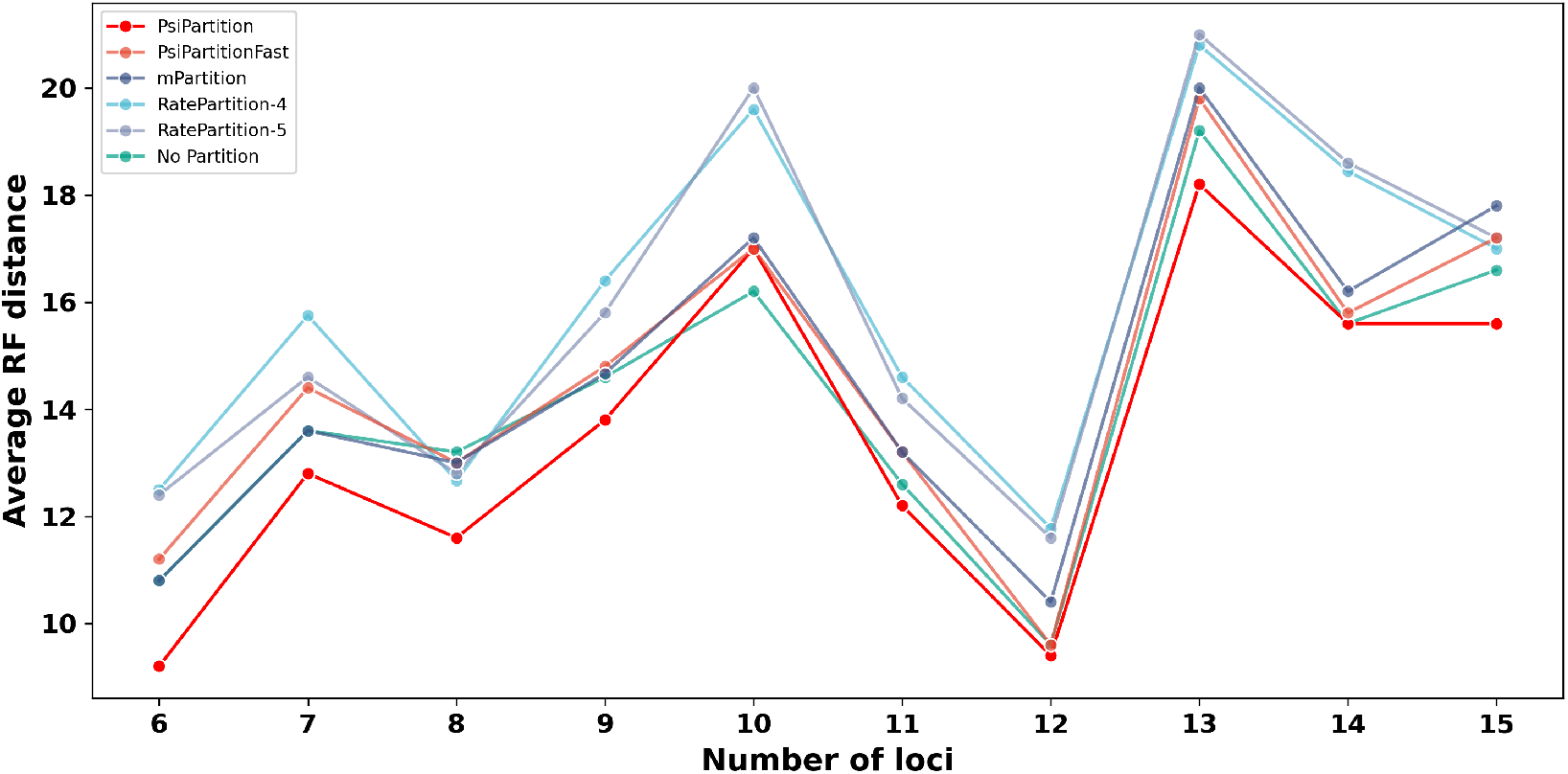
The average RF distances between the true trees and the reconstructed trees on SimDNA-2. PsiPartition (red) reconstructs the most accurate trees with the smallest RF distances for most of number of loci. The only exception when the number of loci is 10 can be explained by the randomness from the relatively small scale of the simulated data set (only 10 trees).

### 3.4. Tree analysis on Noctuidae data set

We reproduced the phylogeny in the reported work [43] by IQ-TREE, and compared it to the phylogeny inferred by PsiPartition. Overall, the results show that the reconstructed tree based on PsiPartition has more likely topology and higher bootstrap supports of the branches, as shown in Supplementary Figure S1, compared to the tree without partitioning in Supplementary Figure S2. To emphasize the accuracy of PsiPartition clearly, we compare the reconstructed topologies and bootstrap supports in detail.

As shown in Fig. 9, in the reproduced phylogeny (tree without partitioning), the species *Arcte modesta* and *Pseudoarcte melanis* are firstly categorized as a clade with a bootstrap support of 57 and then categorized with *Paracte schneideriana* into a clade with a bootstrap support of 100. The reproduced tree has the same topology and similar bootstrap support (65) to the tree in the reported work [43]. Both of them are below the commonly used threshold of 70 and are not convinced. On the other hand, in the tree generated by PsiPartition, *Arcte modesta* and *Paracte schneideriana* are firstly categorized into a clade with a bootstrap support of 93 and then categorized with *Pseudoarcte melanis* into a clade with a bootstrap support of 100, which are more convincing, suggesting a possibly correct reconstruction of these species.

**Fig. 9.**
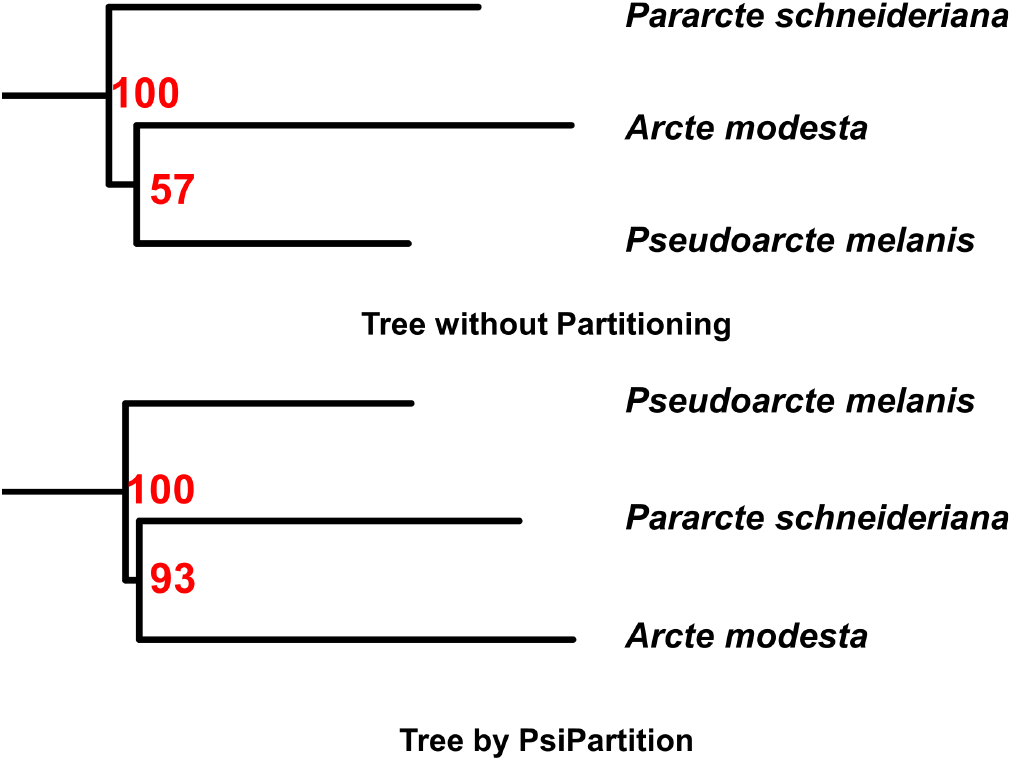
The reconstructed trees of *Arcte modesta, Pseudoarcte melanis*, and *Paracte schneideriana* from phylogeny without partitioning (above) and PsiPartition (below), which shows a possible correct topology with higher bootstrap supports.

As shown in Fig. 10, in the reproduced phylogeny (tree without partitioning), the species *Diloba caeruleocephala, Raphia abrupta or sp, Deltote uncula*, and *Cucullia unmbratica* are nested into three clades with bootstrap supports (43/81/59). In the reported work [43], *Deltote uncula* and *Cucullia unmbratica* are also categorized as a single clade with posterior probabilities of 0.61, which is very low. On the other hand, in the tree generated by PsiPartition, *Deltote uncula, Diloba caeruleocephala, Raphia abrupta or sp*, and *Cucullia umbratica* are nested into three clades with bootstrap supports (90/89/77). Overall, PsiPartition provides a different and possibly more accurate reconstruction of these species compared to the original work and reproduced work and reasons could be discussed in future work.

**Fig. 10.**
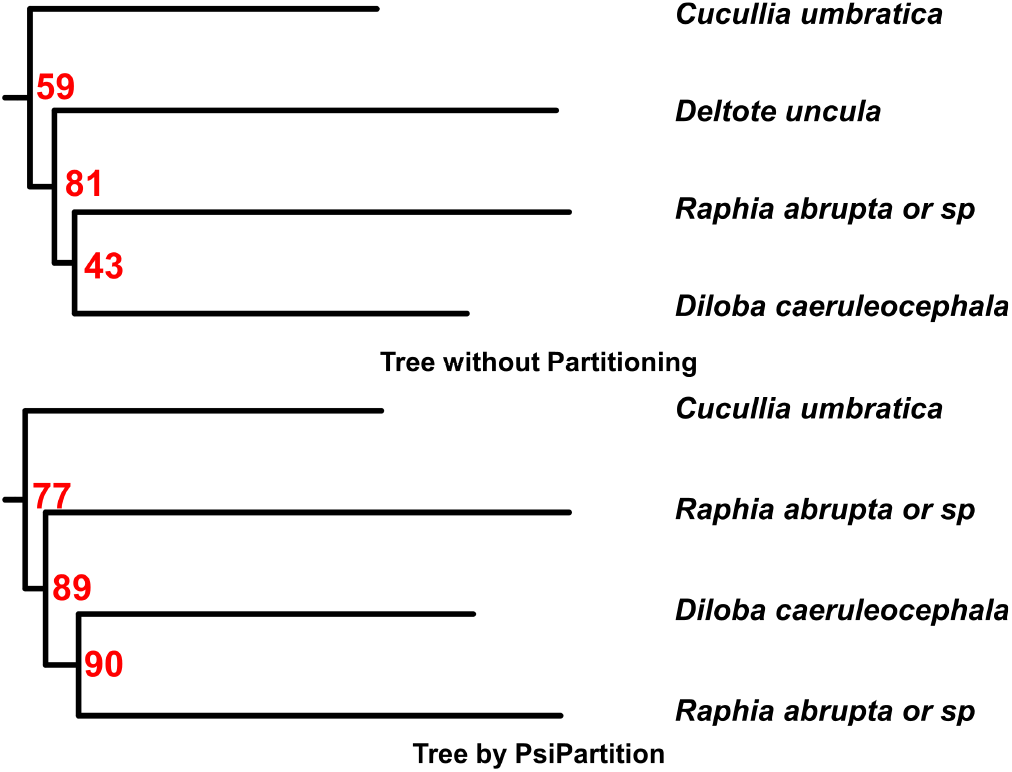
The reconstructed clades of *Diloba caeruleocephala, Raphia abrupta or sp, Deltote uncula*, and *Cucullia unmbratica* from phylogeny without partitioning (above) and PsiPartition (below), which shows a possible correct topology with higher bootstrap supports.

As shown in Fig. 11, both the reproduced phylogeny (tree without partitioning) and the tree generated by PsiPartition reconstructed the same topology for the species of *Craniophora ligustri, Polygrammate hebraeicum, Comachara cadburyi, Harrisimemna trisignata*, and *Cerma cerintha* yet with different bootstrap supports (98/100/62/51 v.s. 100/100/82/80). In the reported work [43], *Craniophora liguistr* together with *Harrisimemna trisignata* and *Cerma cerintha* are categorized into a clade with very low posterior probabilities of 0.59. PsiPartition thus provides an accurate reconstruction of this clade with higher bootstrap supports. These results demonstrate that, by properly partitioning sites with our proposed PsiPartition method, we can improve the accuracy of the reconstructed phylogenetic trees ands the bootstrap supports of the branches.

**Fig. 11.**
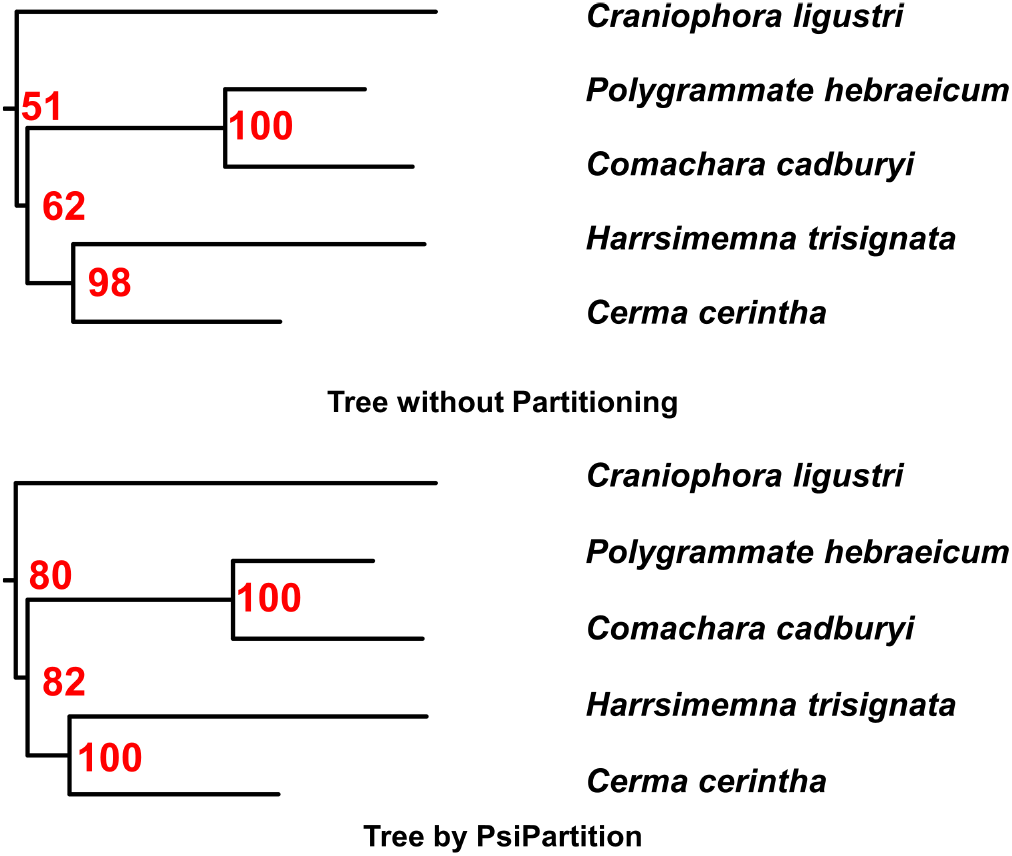
The reconstructed clades of *Craniophora ligustri, Polygrammate hebraeicum, Comachara cadburyi, Harrisimemna trisignata*, and *Cerma cerintha* from phylogeny without partitioning (above) and PsiPartition (below). They have the same topology yet different bootstrap supports on some branches.

## 4. Conclusion

In this study, we proposed a novel partition approach based on parameterized sorting indices (PSI) and Bayesian optimization to accurately and efficiently obtain the optimal partitioning of sites in genomic data. We applied our method to empirical data sets and achieved the best information criterion (BIC and AICc) compared to the models without partitioning or other existing state-of-the-art methods. We also show that Bayesian optimization-based PsiPartition is sufficiently efficient to determine the optimal parameters for partitioning the sites. In addition, we tested PsiPartition on the challenging simulated data sets with various site heterogeneity and demonstrated that our method outperforms other methods in terms of the average RF distance of the reconstructed phylogenetic trees to the actual trees. Finally, we show that on the empirical data set Noctuidae, our proposed method can improve the accuracy of the reconstructed phylogenetic trees, as well as the bootstrap supports of the branches. Overall, our proposed Bayesian optimization-based method achieves the best performance and provides a novel general framework to efficiently determine the optimal number of partitions, which can be easily extended to other partitioning methods. We hope its efficiency and effectiveness contribute to the growing demand for partitioning methods in phylogenetic analysis of large-scale genomic data.

## CRediT authorship contribution statement

**Shijie Xu:** Methodology, Software, Validation, Formal analysis, Investigation, Data curation, Writing Original Draft, Visualization, Writing Review & Editing, Funding acquisition. **Akira Onoda:** Resources, Writing Review & Editing, Supervision, Project administration, Funding acquisition.

## Code availability

The corresponding reproducible source code, model weights, datasets are publicly available at https://github.com/xu-shi-jie/PsiPartition.

## A. Supplementary data

Supplementary data to this article can be found online at https://doi.org/xxxx.

