## Supplementary material for "PsiPartition: Improved Site Partitioning for Genomic Data by Parameterized Sorting Indices and Bayesian Optimization": SI

**Table S1.** Performance comparison of different partitioning schemes on the empirical DNA data sets.

| Data set | LogL | AICc | BIC | AIC | # params | Method |
|---|---|---|---|---|---|---|
| Arctiina | -46408 | 93620 | 96075 | 93568 | 376 | SortPartition |
| Arctiina | -50345 | 101408 | 103619 | 101366 | 338 | RatePartition-4 |
| Arctiina | -51181 | 102857 | 104417 | 102837 | 237 | No Partition |
| Arctiina | -46814 | 94426 | 96862 | 94375 | 373 | SortPartitionFast |
| Arctiina | -50254 | 101274 | 103620 | 101226 | 359 | RatePartition-5 |
| Arctiina | -49255 | 99044 | 100721 | 99021 | 255 | mPartition |
| Calisto | -42277 | 85122 | 86868 | 85093 | 270 | RatePartition-4 |
| Calisto | -43048 | 86492 | 87734 | 86478 | 191 | No Partition |
| Calisto | -38951 | 78671 | 80973 | 78619 | 358 | SortPartition |
| Calisto | -38832 | 78379 | 80536 | 78334 | 335 | SortPartitionFast |
| Calisto | -41304 | 83045 | 84408 | 83027 | 210 | mPartition |
| Calisto | -42266 | 85155 | 87053 | 85120 | 294 | RatePartition-5 |
| Choreutidae | -57403 | 115381 | 117211 | 115356 | 275 | RatePartition-5 |
| Choreutidae | -60854 | 121889 | 122487 | 121886 | 89 | No Partition |
| Choreutidae | -58749 | 117716 | 118434 | 117712 | 107 | mPartition |
| Choreutidae | -55585 | 111529 | 112700 | 111519 | 175 | SortPartitionFast |
| Choreutidae | -54428 | 109383 | 111069 | 109362 | 253 | SortPartition |
| Choreutidae | -57509 | 115505 | 117065 | 115486 | 234 | RatePartition-4 |
| Coenonymphina | -62374 | 125319 | 126999 | 125285 | 268 | RatePartition-5 |
| Coenonymphina | -59165 | 119201 | 121663 | 119123 | 397 | SortPartition |
| Coenonymphina | -62311 | 125148 | 126705 | 125119 | 248 | RatePartition-4 |
| Coenonymphina | -61796 | 123941 | 125002 | 123927 | 168 | mPartition |
| Coenonymphina | -64339 | 128991 | 129946 | 128980 | 151 | No Partition |
| Coenonymphina | -60631 | 121908 | 123790 | 121864 | 301 | SortPartitionFast |
| Geometridae | -188014 | 377098 | 380300 | 377010 | 491 | RatePartition-4 |
| Geometridae | -184958 | 371222 | 375039 | 371093 | 589 | SortPartitionFast |
| Geometridae | -191587 | 383950 | 386341 | 383903 | 364 | mPartition |
| Geometridae | -192079 | 384890 | 387159 | 384848 | 345 | No Partition |
| Geometridae | -184394 | 370294 | 374608 | 370126 | 669 | SortPartition |
| Geometridae | -188009 | 377145 | 380498 | 377048 | 515 | RatePartition-5 |
| Morpho | -28203 | 56577 | 57143 | 56575 | 84 | mPartition |
| Morpho | -24007 | 48397 | 49643 | 48386 | 186 | SortPartition |
| Morpho | -27741 | 55862 | 57095 | 55851 | 184 | RatePartition-4 |
| Morpho | -29365 | 58873 | 59351 | 58871 | 71 | No Partition |
| Morpho | -25705 | 51691 | 52611 | 51684 | 137 | SortPartitionFast |
| Morpho | -27667 | 55756 | 57122 | 55743 | 204 | RatePartition-5 |
| Noctuidae | -96323 | 193446 | 195940 | 193399 | 376 | SortPartition |
| Noctuidae | -100391 | 201148 | 202340 | 201137 | 178 | mPartition |
| Noctuidae | -101258 | 203271 | 205635 | 203229 | 356 | RatePartition-4 |
| Noctuidae | -101074 | 203032 | 205766 | 202974 | 413 | RatePartition-5 |
| Noctuidae | -103295 | 206937 | 208070 | 206927 | 169 | No Partition |
| Noctuidae | -97557 | 195752 | 197776 | 195721 | 304 | SortPartitionFast |
| Pieridae | -135110 | 271296 | 274547 | 271210 | 495 | RatePartition-5 |
| Pieridae | -131424 | 263804 | 266728 | 263736 | 444 | SortPartitionFast |
| Pieridae | -135102 | 271187 | 274182 | 271115 | 455 | RatePartition-4 |
| Pieridae | -138384 | 277253 | 278805 | 277235 | 233 | No Partition |
| Pieridae | -136377 | 273258 | 274870 | 273239 | 242 | mPartition |
| Pieridae | -131295 | 263477 | 266208 | 263418 | 414 | SortPartition |

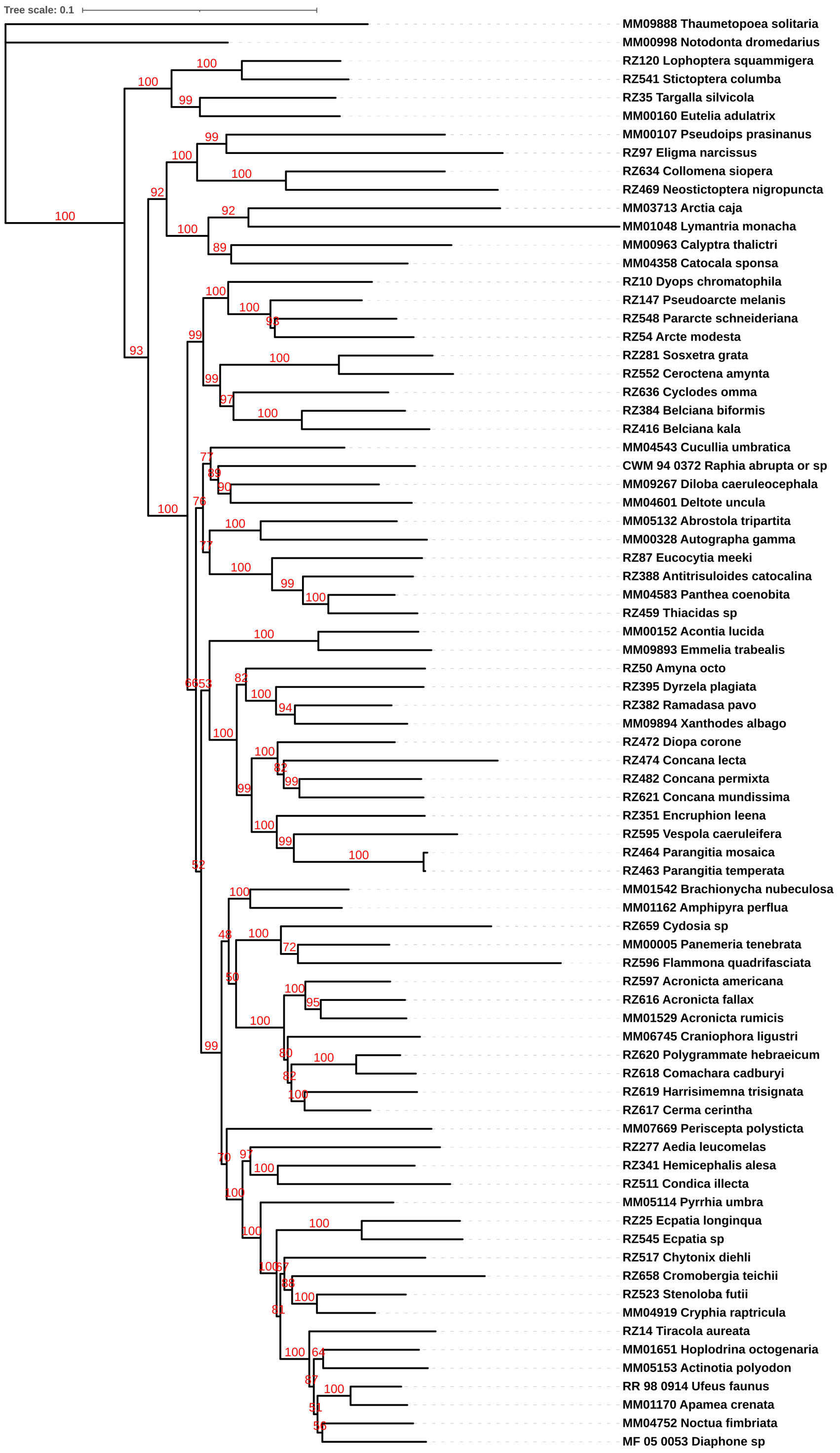


**Figure S1.** The reconstructed Noctuidae tree based on PsiPartition.

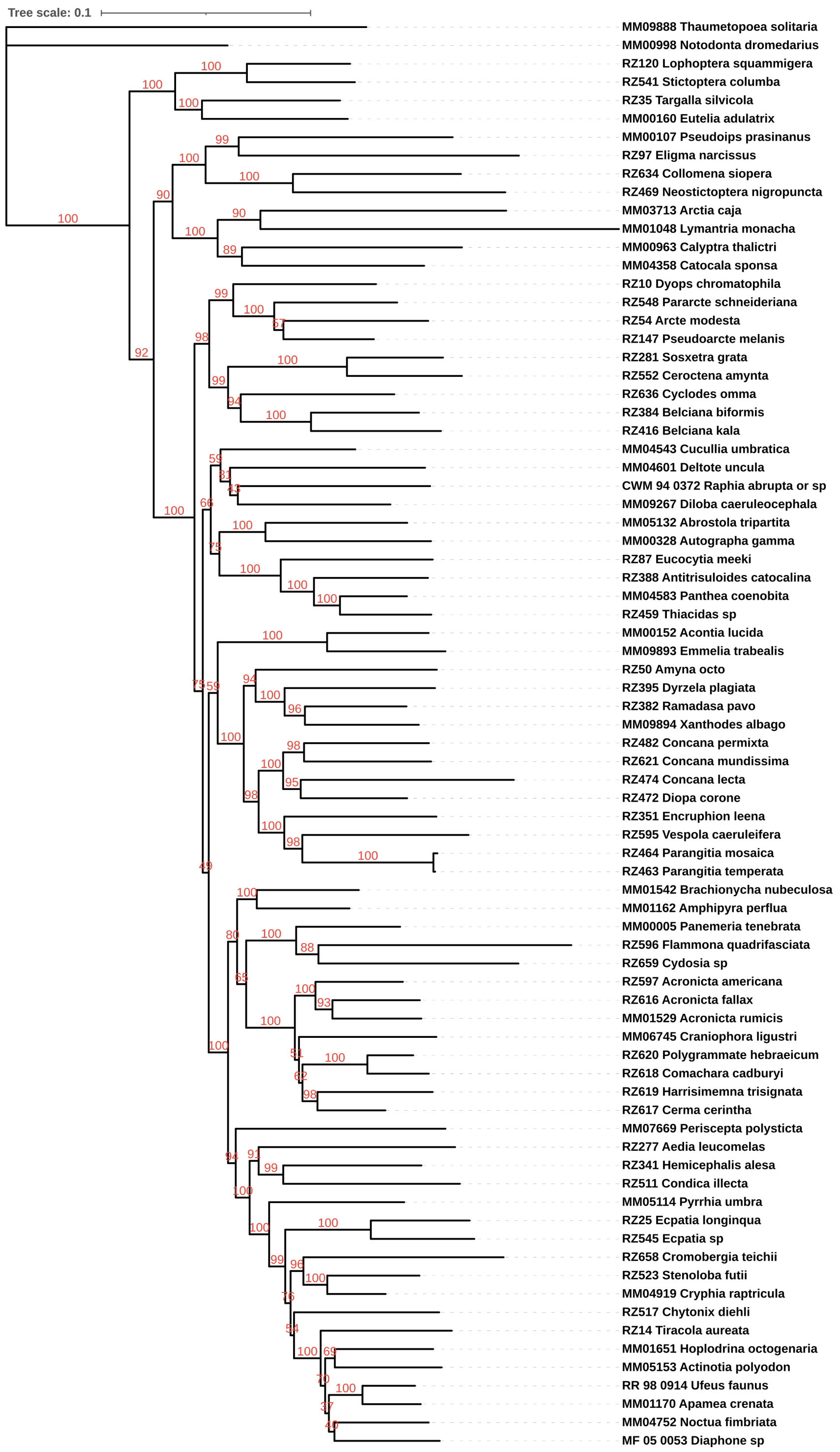


**Figure S2.** The reconstructed Noctuidae tree without partitioning.

**Algorithm S1** PsiPartitionFast

**Require:**

$A$: the alignment
$k_{max}$: the maximum number of partitions
$n$: the maximum iterations steps

**Ensure:**

$P^*$: the optimal partition
$s^*$: the optimal parameters
$o^*$: the optimal objective value

Uniformly sample $k$ from $2, 3, ..., k_{max}$
Uniformly sample $w$ from $[0, 1]^t$ ▷ $t = 5$ for DNA, 21 for protein
Let $S$ be an empty set
Initialize a surrogate model $m$
**for** $i = 1$ to $n$ **do**
    Let $\Psi$ be the PSI of all sites with parameters $w$
    Bin the sites of $A$ into $k$ partitions based on $\Psi$ as $P_i$
    Evaluate IQ-TREE with partitioning $P_i$ and obtain BIC $b_i$
    $s_i \leftarrow k$
    $S \leftarrow S \cup \{(P_i, s_i, b_i)\}$
    Update $m$ with $S$
    Determine the next parameters $s_{i+1}$ by the surrogate model $m$ with expected improvement
**end for**
return $(P^*, s^*, o^*) \in S$ that has the minimum BIC $o^*$

**Algorithm S2** Generate binary tree

**Require:**

$n$: the number of leaves

**Ensure:**

$T$: a binary tree with $n$ leaves

Let $S$ be a set of $n$ nodes
$i \leftarrow n$
**while** $|S| > 1$ **do**
    Randomly choose two nodes $u$ and $v$ from $S$
    Construct a tree $T$ with $u$ and $v$ as its children
    Remove $u$ and $v$ from $S$ and add $T$ to $S$
**end while**
return the only tree in $S$

**Table S2.** Performance comparison of different partitioning schemes on the empirical protein data sets.

| Data set | LogL | AICc | BIC | AIC | # params | Method |
|---|---|---|---|---|---|---|
| Nguyen_2016a | -3979 | 8041 | 8213 | 8037 | 39 | RatePartition-4 |
| Nguyen_2016a | -3138 | 6674 | 7282 | 6584 | 154 | SortPartition |
| Nguyen_2016a | -3944 | 7971 | 8143 | 7966 | 39 | RatePartition-5 |
| Nguyen_2016a | -4270 | 8609 | 8755 | 8606 | 33 | No Partition |
| Nguyen_2016a | -3858 | 7841 | 8089 | 7830 | 57 | SortPartitionFast |
| Nguyen_2016a | -4139 | 8357 | 8520 | 8352 | 37 | mPartition |
| Nguyen_2016b | -5370 | 10882 | 11158 | 10868 | 64 | mPartition |
| Nguyen_2016b | -5541 | 11216 | 11479 | 11203 | 61 | No Partition |
| Nguyen_2016b | -5153 | 10496 | 10848 | 10472 | 83 | SortPartitionFast |
| Nguyen_2016b | -5059 | 10268 | 10556 | 10253 | 67 | RatePartition-4 |
| Nguyen_2016b | -4599 | 9712 | 10412 | 9571 | 186 | SortPartition |
| Nguyen_2016b | -5057 | 10269 | 10565 | 10253 | 69 | RatePartition-5 |
| Nguyen_2016c | -5874 | 11909 | 12238 | 11895 | 73 | SortPartitionFast |
| Nguyen_2016c | -6298 | 12749 | 13065 | 12736 | 70 | No Partition |
| Nguyen_2016c | -5223 | 10899 | 11631 | 10799 | 177 | SortPartition |
| Nguyen_2016c | -5936 | 11995 | 12254 | 11986 | 57 | RatePartition-4 |
| Nguyen_2016c | -5930 | 12030 | 12375 | 12014 | 77 | RatePartition-5 |
| Nguyen_2016c | -6104 | 12363 | 12683 | 12349 | 71 | mPartition |
| Nguyen_2016d | -5845 | 11817 | 12071 | 11807 | 58 | SortPartitionFast |
| Nguyen_2016d | -5343 | 11119 | 11769 | 11017 | 165 | SortPartition |
| Nguyen_2016d | -5804 | 11752 | 12034 | 11738 | 65 | RatePartition-5 |
| Nguyen_2016d | -5833 | 11760 | 11955 | 11754 | 44 | RatePartition-4 |
| Nguyen_2016d | -6087 | 12247 | 12403 | 12243 | 35 | mPartition |
| Nguyen_2016d | -6312 | 12698 | 12854 | 12694 | 35 | No Partition |
| Nguyen_2016e | -3388 | 6921 | 7139 | 6897 | 61 | SortPartition |
| Nguyen_2016e | -3399 | 6895 | 7054 | 6884 | 43 | RatePartition-4 |
| Nguyen_2016e | -3592 | 7262 | 7393 | 7255 | 35 | mPartition |
| Nguyen_2016e | -3438 | 7013 | 7221 | 6992 | 58 | SortPartitionFast |
| Nguyen_2016e | -3686 | 7441 | 7562 | 7435 | 32 | No Partition |
| Nguyen_2016e | -3397 | 6896 | 7061 | 6883 | 45 | RatePartition-5 |
| Nguyen_2016f | -4000 | 8137 | 8389 | 8122 | 61 | SortPartitionFast |
| Nguyen_2016f | -3983 | 8160 | 8495 | 8132 | 83 | RatePartition-5 |
| Nguyen_2016f | -3997 | 8133 | 8388 | 8118 | 62 | RatePartition-4 |
| Nguyen_2016f | -3378 | 7186 | 7755 | 7069 | 157 | SortPartition |
| Nguyen_2016f | -4437 | 8944 | 9084 | 8940 | 33 | No Partition |
| Nguyen_2016f | -4288 | 8651 | 8800 | 8647 | 35 | mPartition |
